# Hyperalignment Reveals Shared Information in Idiosyncratic Fine-Grained Movie fMRI Patterns in Macaques

**DOI:** 10.64898/2026.06.14.731832

**Authors:** Yuqi Zhang, Haiyan Wang, Qi Zhu, Xiaolian Li, Jane Han, Guo Jiahui, Won Mok Shim, Royoung Kim, Maria Ida Gobbini, Kwon Heo, Alessia Sepe, James V. Haxby, Ma Feilong, Wim Vanduffel

**Affiliations:** Department of Psychological and Brain Sciences, Dartmouth College, Hanover, NH, USA; Beijing Key Laboratory of Brainnetome and Brain-Computer Interface, Institute of Automation, Chinese Academy of Sciences, Beijing, China; Brainnetome Center, Institute of Automation, Chinese Academy of Sciences, Beijing, China; Laboratory of Neuro- and Psychophysiology, Department of Neurosciences, KU Leuven Medical School, Leuven, Belgium; Leuven Brain Institute, KU Leuven, Leuven, Belgium; Cognitive Neuroimaging Unit, INSERM, CEA, Université Paris-Saclay, NeuroSpin Center, Gif-sur-Yvette, France; Department of Psychology, School of Behavioral and Brain Sciences, University of Texas at Dallas, Richardson, TX, USA; Center for Vital Longevity, School of Behavioral and Brain Sciences, University of Texas at Dallas, Dallas, TX, USA; Department of Intelligent Precision Healthcare Convergence, Sungkyunkwan University, Suwon, South Korea; Center for Neuroscience Imaging Research, Sungkyunkwan University, Suwon, South Korea; Department of Biomedical Engineering, Sungkyunkwan University, Suwon, South Korea; Department of Medical and Surgical Sciences, University of Bologna, Bologna, Italy; IRCCS Institute of Neurological Sciences of Bologna, Bologna, Italy; Department of Psychology, University of South Carolina, Columbia, SC, USA; Institute for Mind and Brain, University of South Carolina, Columbia, SC, USA; Athinoula A. Martinos Center for Biomedical Imaging, Massachusetts General Hospital, Charlestown, MA, USA; Department of Radiology, Harvard Medical School, Boston, MA, USA

## Abstract

Response hyperalignment (RHA) of fMRI patterns acquired during movie-viewing in 12 rhesus monkeys revealed shared representational spaces despite idiosyncratic functional topographies. RHA greatly improved inter-subject correlation of movie response time-series in occipital, temporal, and prefrontal cortices and afforded highly accurate between-subject decoding of movie time-points and estimation of idiosyncratic functional topographies. These results demonstrate that nonhuman primates share high-dimensional, high-capacity representational spaces encoded in idiosyncratic fine-grained functional topographies.

## Main

How shared neural representations are encoded despite substantial inter-individual variability in the functional organization of the cortex remains unclear. Here we applied response hyperalignment (RHA) to fMRI patterns acquired during naturalistic movie viewing in macaques to model shared representational spaces encoded in fine-scale response patterns and to predict idiosyncratic coarse-scale category-selective topographies.

Information is encoded in neural population responses, and these representations can be decoded in monkeys from recordings of multiple single units with pattern classifiers^1,2^ or representational similarity analysis^3,4^. Similarly, population responses also can be decoded from fMRI data in both humans^5,6^ and monkeys^7–10^, indicating that the underlying neuronal tuning profiles have a fine-grained topographic structure. This fine-scale functional organization affords decoding of response patterns within category-selective areas, such as the monkey face and body patches^11,12^, or human face and place areas^5,13–15^. In parallel, cortical functional topography also has a coarse-scale structure that separates different sensory, motor, and cognitive functions into large areas, pathways, and systems^16,17^. Unlike this coarse-scale organization, fine-grained topographies cannot be aligned across brains using anatomical features. Consequently, decoding is usually done within individual brains, and results are aggregated across brains as mean accuracies. Whether the neuronal population codes that underlie accurate decoding in monkeys share a common representational structure across individuals therefore remains unknown.

RHA models the shared information embedded in idiosyncratic fine-scale functional topographies^14,18–21^. It first derives a common model space that has an anatomical structure in which each common model dimension is a location in a high-resolution cortical model. This common space is estimated from neural response patterns to naturalistic stimuli (usually movies), capturing representations of a broad range of sensory inputs and cognitive processes. In a second step, a subject-specific transformation matrix is calculated to project idiosyncratic individual cortical vertex spaces into the common model space. These matrices can then be used to project new data into that common space.

RHA is a flexible computational framework originally developed to model fine-scale functional topographies in human fMRI data and has much broader applications. Resampling vertex time-series responses into the common model space greatly improves inter-subject correlations of responses and connectivity profiles, thereby enhancing between-subject decoding accuracy. RHA has also been applied to single-unit recordings from the rat hippocampus to derive a common representational space of navigation^22^. Interestingly, RHA also afforded high-fidelity modeling of coarse-scale category-selective and retinotopic topographies by projecting fMRI localizer data from multiple brains into an index brain^18,19,23,24^.

We applied response RHA to fMRI data collected in 12 macaques during naturalistic movie viewing of the commercial movie, *Monkey Kingdom*. fMRI responses to the first three clips of the movie were used as training data to construct a common representational space and estimate subject-specific transformations that aligned individual brains into that space. These transformations were then applied to hyperalign responses of the left-out two clips of the movie (test data) to evaluate whether RHA improved inter-subject correlation of movie-evoked responses, enhanced between-subject decoding of movie time-points, and enabled prediction of individual functional topographies from functional localizer data acquired in other monkeys.

Compared to anatomical alignment, RHA increased inter-subject correlation (ISC) across most of the cortex (Fig. 1a). Mean ISC across all cortical vertices nearly doubled (from 0.253 to 0.496). The strongest gains were observed in occipital, temporal, and lateral prefrontal cortices (Fig. 1a), showing that RHA improved alignment not only in visual areas but also in higher order cortical systems. All 12 macaques showed a large increase of ISC with RHA (Fig. 1b), indicating that the improvement was consistent across macaques rather than driven by few subjects.

**Figure 1:**
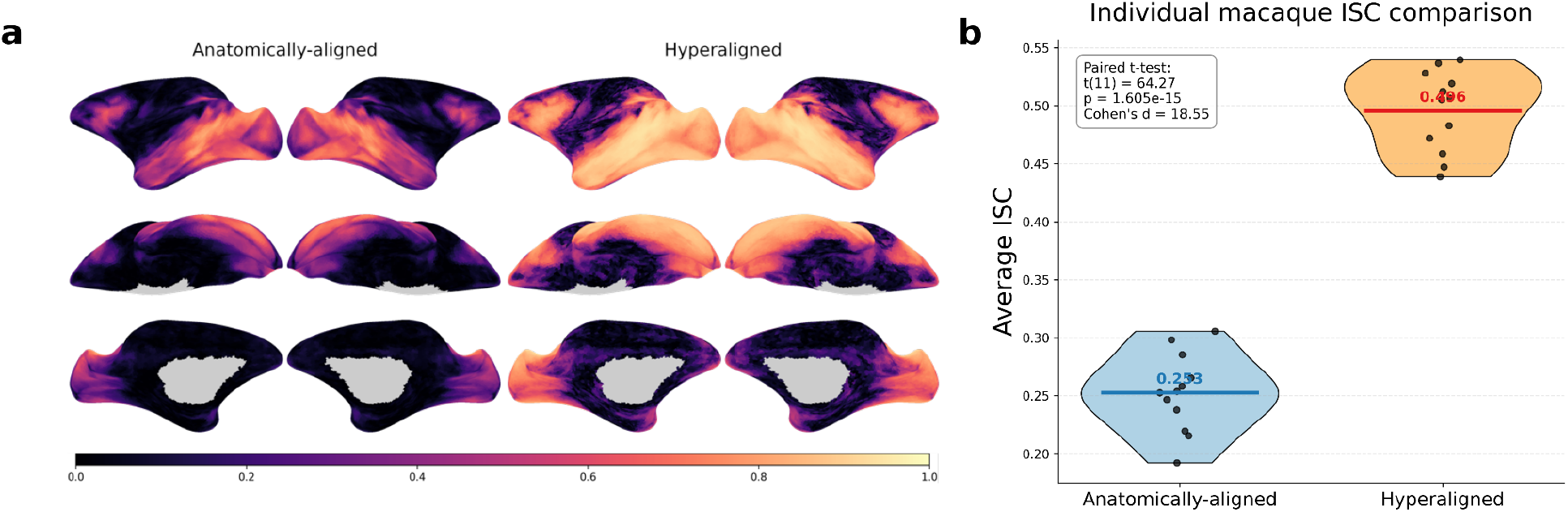
RHA increases inter-subject correlation of movie-evoked responses in macaques. (a) Group-average inter-subject correlation (ISC) maps for anatomically-aligned and hyperaligned macaque data during movie viewing. (b) Distribution of average whole-brain ISC values across 12 macaques for anatomically-aligned and hyperaligned data. Each dot represents one macaque. Mean ISC increased from 0.253 under anatomical alignment to 0.496 after RHA (paired *t*-test *t*(11) = 64.27, p = 1.605×10^−15^, Cohen’s *d* = 18.55).

We next tested whether we could decode which part of the movie a macaque was watching by comparing the individual response patterns to the mean patterns in other macaques. We performed between-subject classification of 2 s long time-points using both whole-brain and searchlight patterns, and nearest-neighbor classification based on correlation distance. For a correct classification, the correlation between an individual macaque’s pattern for a single time-point with the mean response pattern to the same time-point in other macaques must be higher than correlations between that time-point in the index macaque and all other time-points in the other macaques. Chance performance is 1/900 (<0.0012).

For whole-brain decoding, we calculated the principal components of response patterns and used the first 10 to 300 principal components. Peak classification accuracy increased by roughly 80%, from 0.240 with anatomical alignment to 0.430 after RHA (Fig. 2a). Over 70 components were required to reach peak performance, and RHA consistently outperformed accuracies based on anatomically-aligned data (Fig. 2b). Restricting the analysis to local response patterns within 8 mm searchlights showed even larger gains, with mean searchlight decoding accuracy increasing more than fivefold (from 0.005 to 0.028 after RHA), with the highest decoding accuracies (0.162) observed in the posterior inferotemporal (IT) cortex (Fig. 2c). These high levels of decoding accuracy indicate that RHA aligns fine-grained cortical response patterns in macaques carrying information that is shared across individuals, with the strongest shared representations in IT and lateral prefrontal cortices.

**Figure 2:**
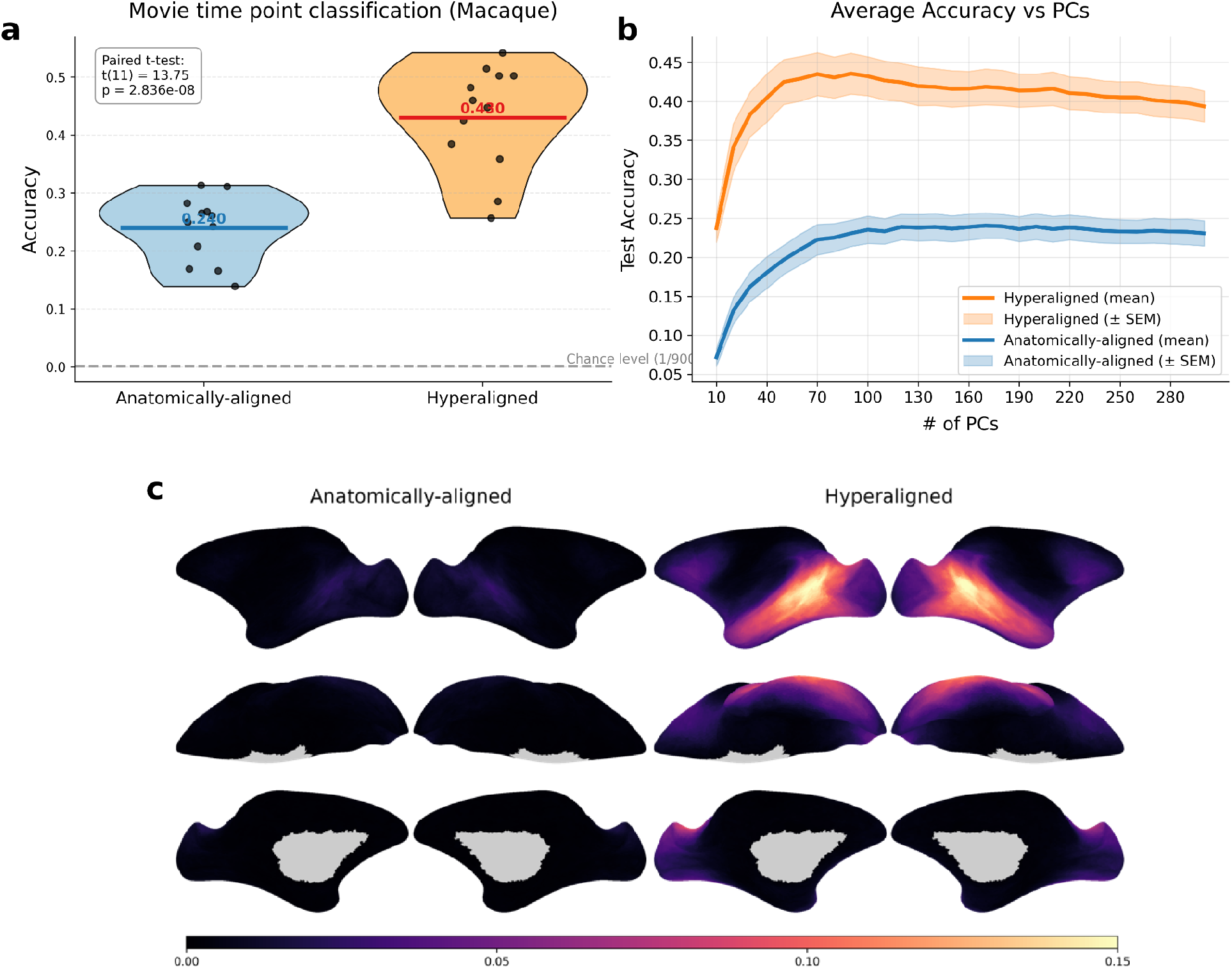
Hyperalignment improves between-subject movie time-point classification in macaques. (a) Whole-brain classification accuracy for movie time-points following anatomically-aligned and RHA in macaques. Each point denotes one macaque, and horizontal lines mark group means. RHA increased mean accuracy from 0.240 to 0.430 (paired *t*-test: *t*(11) = 13.75, p = 2.836×10^−8^). The dashed line shows chance performance (1/900). (b) Average classification accuracy as a function of the number of principal components. Shaded regions denote ± SEM. (c) Searchlight-based classification accuracy maps for anatomically- and hyperaligned data.

Finally, we tested whether RHA could predict coarse-scale individual functional topographies from other macaque data. Visual category localizer data were available for four macaques from independent studies using still images of faces, bodies, and objects^15,25,26^. We used pairwise hyperalignment transforms estimated from movie responses to project category-selective localizer maps (face-versus-object and bodies-versus-object) from the other macaques into the cortical anatomy of each of these four macaques.

Although macaques showed broadly similar category-selective topographies, each individual showed a distinct idiosyncratic functional organization (Fig. 3a, left). The predicted maps accurately recovered these individual-specific topographies (Fig. 3a, right), including canonical face-patches such as ML/MF and AL^8^ (Supp. Fig. S1). To quantify recovery of individual-specific topographies, we calculated correlations between each individual’s measured and predicted category-selective maps within IT cortex (congruent maps) and compared these to correlations between each individual’s maps and the predicted maps of other individuals (incongruent maps). For each macaque and both category-selective maps, congruent predictions consistently exceeded incongruent ones (Fig. 3b). These results show that the RHA transformation matrices, based on responses to a naturalistic movie, accurately capture individual differences in coarse-scale category-selective functional topography.

**Figure 3:**
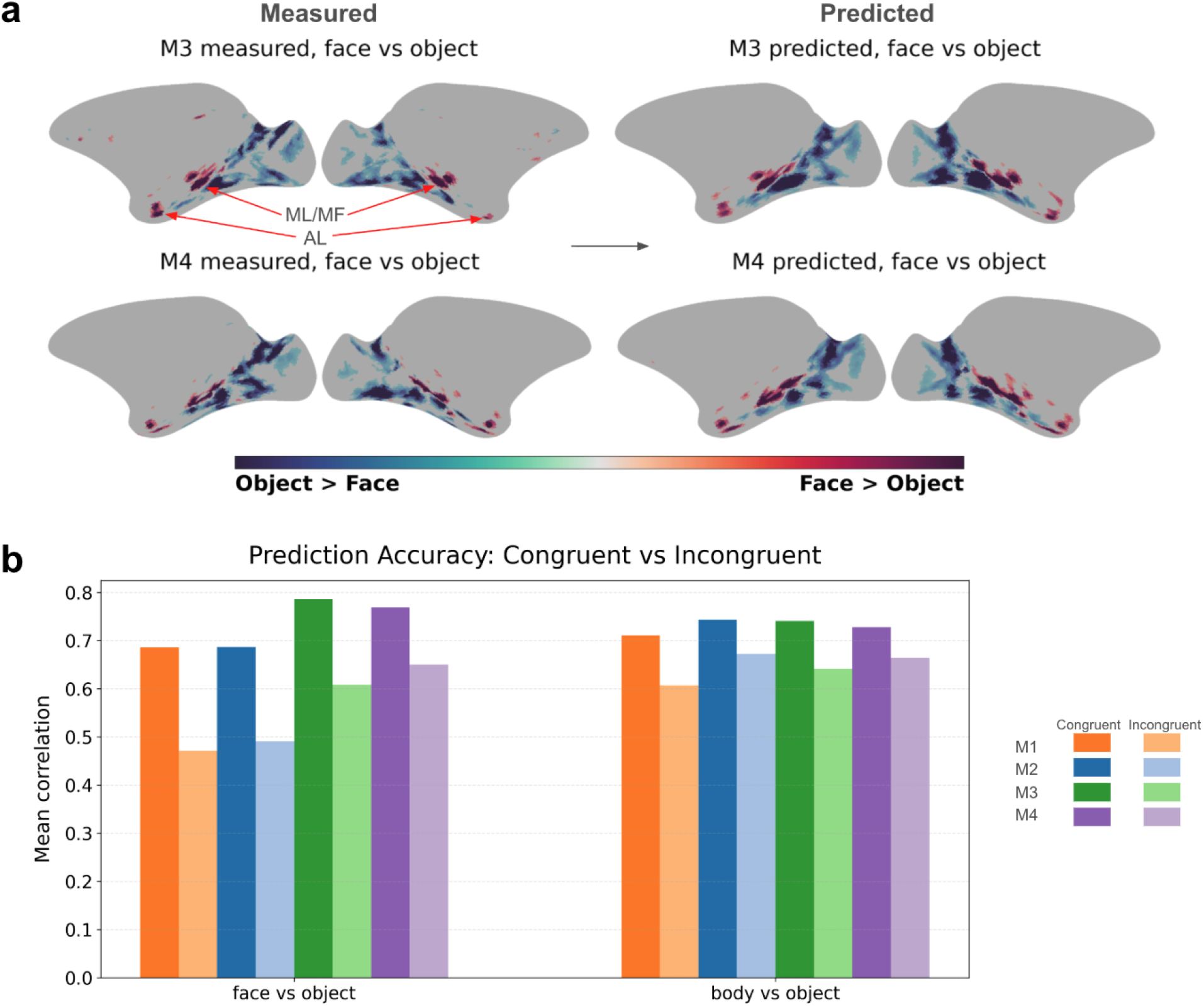
Hyperalignment enables prediction of individual functional topographies in macaques. (a) Measured and predicted (z-scored) face-versus-object localizer maps, thresholded to retain only the upper and lower 5% of z-score values, are shown for two example macaques (M3 and M4). Predicted maps recovered subject-specific features, including face-selective patches such as ML/MF and AL. (b) Mean correlations between measured patterns and congruent or incongruent predicted patterns for face-versus-object and body-versus-object localizers across four macaques.

Naturalistic stimulus paradigms, such as movie-viewing, afford sampling a broad range of stimuli and cognitive processing, including dynamic agentic and social actions^21,27–29^, and are critical for deriving a common model of representational spaces that have general validity^14^.

Between-subject classification of movie time-points shows that representational spaces in macaque brains contain hundreds of shared, distinct representations of these dynamic, naturalistic stimuli. Maximal classification performance, based on whole-brain patterns, required more than 70 orthogonal dimensions (Fig. 2b), indicating that simple representational models based on only a few category contrasts are insufficient to model this representational space. In addition to modeling shared information encoded in fine-scale topographies, RHA of macaque movie data also captured individual differences in coarser-scale category-selective functional topographies. RHA was also data-efficient, as transformations estimated from only two movie repetitions produced higher ISC than anatomical alignment using seven repetitions, and ISC continued to improve with additional hyperaligned data (Supp. Fig. S7).

In sum, our results show that shared information in high-dimensional and high-capacity cortical representational spaces in the macaque brain can be measured with fMRI during naturalistic stimulation and modeled with RHA. These representational spaces capture both shared information encoded in idiosyncratic fine-scale topographies and individual differences in coarser-scale functional topographies. Similar results have been reported in human fMRI studies using hyperalignment^14,18,19,21,23,24^, and parallel human analyses showed the same overall pattern using data obtained from humans watching the same movie and performing object selective tasks (see Supp. Figs. S2-S4). Together, these findings indicate that nonhuman and human primates both have high-dimensional and high-capacity cortical representational spaces that can be revealed with naturalistic fMRI and modeled with hyperalignment.

## Methods

### Subjects

Twelve rhesus monkeys (Macaca mulatta; 1 female; 5.15~9.30 kg) were used for the fMRI study. Animal care and experimental procedures were performed in accordance with the National Institutes of Health’s Guide for the Care and Use of Laboratory Animals, the European legislation (Directive 2010/63/EU) and were approved by the Animal Ethics Committee of the KU Leuven. The Weatherall Report was used as reference for animal housing and handling. All animals were group-housed in cages sized 16~32 m^3^, which encourages social interactions and locomotor behavior. The environment was enriched by foraging devices and toys. The animals were fed daily with standard primate chow supplemented with fruits, vegetables, bread, peanuts, cashew nuts, raisins and dry apricots. The animals were exposed to natural light and additional artificial light for 12 hours every day. On training and experimental days, the animals were allowed unlimited access to fluid through their performance during the experiments. Using operant conditioning techniques with positive reinforcers, the animals received fluid rewards for every correctly performed trial. During non-working days, they received water in their living quarters.

### Datasets & Preprocessing

Twelve macaques were scanned at KU Leuven using a 3T Siemens PRISMA scanner. For movie fMRI data, ten animals were scanned at the spatial resolution of 1.25 × 1.25 × 1.25 mm^3^ using external phased-array receive coils, whereas two were scanned at the spatial resolution of 0.8 × 0.8 × 0.8 mm^3^ using phased-array receive coils embedded in MRI-compatible headposts above the skull. The repetition time (TR) was 1 s for eight monkeys and 2 s for the remaining four monkeys. To enhance the contrast-to-noise ratio, macaques were administered 8~11 mg/kg monocrystalline iron oxide nanoparticles (MION, BioPAL) immediately prior to scanning^30^. Subjects were trained by operant conditioning to perform a free-viewing task using liquid rewards (juice droplets). The movie clips were projected using a Barco LCD projector at 60 Hz refresh rate and 1400 × 1050 resolution onto a translucent screen located at 57 cm from the monkeys’ eyes. A virtual fixation window (21 × 12 degrees) spanning the entire screen was defined, and subjects were trained to maintain their gaze within this window for extended periods. The auditory stimuli were presented through MR-compatible headphones^31^, customized for monkeys (MR Confon GmbH, Magdeburg, Germany). Seven of the monkeys were also trained to keep both hands stationary on response buttons inside a response box. Juice rewards were delivered at variable intervals (800~2,500 ms), contingent on viewing the movie and maintaining hand position. All sessions were conducted with the subject in a head-fixed sphinx position. A photocell, mounted on the bottom-right corner of the monitor and invisible to the subjects, marked successive 2 s movie epochs to synchronize behavioral and neural recordings. Eye position was monitored at 120 Hz using an infrared eye-tracking system (ISCAN, Woburn, MA, USA). All macaques watched the movie *Monkey Kingdom* in five continuous clips in Spanish (940 seconds per clip), with each macaque viewing the movie 7~11 times in the scanner. Fixation within the window for each run was 89.8~100% (mean = 98.33, std = 1.77). The first 40 s (10 s static image, 30 s overlap with the previous movie clip) fMRI signal was removed from the study. To match the temporal resolution across datasets, the fMRI time-series for each clip were resampled to 450 time-points. Face- and body-selective localizer data were also collected for 4 of the 12 macaques^15,26,32^.

Macaque cortical surface reconstruction was performed using a revised version of the NHP pipeline^33^, based on high-resolution T1-weighted (0.4 mm) and, when available, T2-weighted (0.4 mm) anatomical images. Initial brain, white-matter and CSF masks were generated using nBEST^34^ and then manually refined. Macaque fMRI data were preprocessed using in-house scripts, including skull stripping, slice-timing correction, motion correction^35^. To reduce frame-to-frame distortions, slice-wise nonlinear correction was performed using a B-spline grid-based registration method^36^. A session-specific field map was used to correct susceptibility-induced EPI distortions. To facilitate functional-to-anatomical registration, a T1-weighted 3D gradient-echo image (GRE) was acquired during the same session as the functional data. For some monkeys, a low-resolution T1-weighted 3D MPRAGE image was also acquired close in time to the fMRI session, providing an intermediate anatomical reference for functional-to-anatomical registration. Candidate registration solutions were generated using registration methods from FSL, Freesurfer, JIP, and ANTs, either individually or in combination. The 3D GRE and low-resolution T1 brain masks were manually refined when necessary to improve registration accuracy. All registrations were visually inspected in multiple anatomical planes, and the final solution was selected according to the alignment performance. The preprocessed macaque fMRI data were first projected onto each subject’s native cortical surface, and then mapped onto the mkavg-ico32 cortical surface template, which was generated from the cortical surfaces of all 12 macaques.

24 humans were scanned at Dartmouth using a 3T Siemens scanner equipped with a 32-channel phased-array head coil. Functional BOLD images were acquired at a spatial resolution of 2.5 × 2.5 × 2.5 mm^3^ with a volume size of 96 × 96 × 52 and TR = 1 s. All subjects watched the movie *Monkey Kingdom* in 5 continuous clips (955 seconds per clip) once outside the scanner with English audio to ensure that they understood the storyline and once inside the scanner with Spanish audio to reduce the influence of language comprehension on cross-species comparisons. For each clip, the fMRI signal corresponding to the first 40 s (10 s static image, 30 s overlap with the previous movie clip) and the last 15 s (a buffer for hemodynamic lag with no stimuli presented) was excluded from the subsequent analyses. Human fMRI data were preprocessed using fMRIPrep 20.2.7^37^ and resampled to the onavg-ico32 cortical surface template^38^.

The same denoising procedure was applied to both human and macaque fMRI data. Nuisance regressors included six head-motion parameters and their temporal derivatives, framewise displacement, 6 principal components extracted from a combined cerebrospinal fluid and white matter mask (aCompCor), and polynomial trends up to the second order.

### Response Hyperalignment

We used response hyperalignment (RHA) to model shared information by projecting individual brain responses into a common model space using movie-viewing data. Whole-brain templates for both macaques and humans were built from the first 3 movie clips using a searchlight-based algorithm that finds shared information via a singular value decomposition of the group data, followed by rotating that information with the Procrustes transformation into the central tendency of topographic distribution (Feilong et al., 2023). There were 19,863 overlapping macaque searchlights (mean size: 167 vertices) and 19,341 human searchlights (mean size: 121 vertices). For each subject, a transformation matrix was estimated with ridge regression using the same three clips and applied to the two held-out clips to project data into the common space. To evaluate RHA performance, we performed two analyses on anatomically-aligned and hyperaligned test movie-viewing data: inter-subject correlation (ISC) of response time-series and movie time-point prediction. We further tested functional map prediction by calculating pairwise RHA transformation matrices from the four macaques for which we obtained localizer data. We used these transformations to project category-selective localizer data into the anatomical space of each target monkey.

### Inter-subject Correlation

To evaluate the cross-subject similarity in movie-evoked responses, we computed vertex-wise inter-subject correlation (ISC). For each vertex, we calculated the Pearson correlation between the response time-series of each subject and the average time-series of all other subjects, where higher ISC indicates better performance. ISC was computed for both anatomically-aligned data and hyperaligned data and then compared across the cortex.

### Movie time-point prediction

Using leave-one-subject-out cross-validation, we performed whole-brain movie time-point prediction in two steps. First, we applied Principal Component Analysis (PCA) to the training subjects to reduce dimensionality. Second, we predicted each time-point in the left-out subject as the nearest-neighbor mean response pattern in the training subjects. The optimal number of principal components was determined from the training data as the value with the highest classification accuracy, and was then used to predict time-points in the left-out test subject.

To determine which brain regions contributed most to prediction performance, we also ran searchlight-based classification using the same searchlight definitions as in the RHA analysis. Time-point classification was computed separately within each searchlight and then averaged across subjects. Accuracy for each searchlight was assigned to its center vertex, to produce whole-brain maps of mean prediction accuracy.

### Functional map prediction

For each of the 4 macaques with visual category localizer data, we computed pairwise RHA transforms from each of the other 3 macaques to the target macaque using all 5 movie clips. We then projected each of the other 3 macaque’s localizer data into the target macaque’s space to generate predicted category-selective maps and averaged the three predictions. Prediction accuracy was quantified as the correlation between each macaque’s measured and predicted localizer maps. To assess individual specificity, we compared this congruent correlation with incongruent correlations, defined as the mean correlation between a macaque’s measured map and the predicted maps from the other macaques.

### Movie repetition analysis

To assess how the amount of repeated movie-viewing data affected alignment performance, we independently varied the number of repetitions used to estimate the transformation and the number used to evaluate it. For each clip, responses were averaged across the first k usable repetitions (k = 1~7, the maximum available across all subjects and clips after quality-control exclusion), concatenated across clips, and z-scored. We used Cronbach’s alpha as the quality-control criterion: If Cronbach’s alpha decreases after including a certain repetition, it means the repetition is likely driven by noise and not used for averaging.

Transformations were estimated from the first three clips using r = 1~7 repetitions. For each repetition count, group templates and subject-specific transformations were estimated using the searchlight procedure described above. Responses to the held-out two clips were independently averaged across d = 1~7 repetitions and projected through the corresponding transformations.

For each combination of transformation and test repetition counts, leave-one-out ISC was calculated at each cortical vertex and averaged across the cortex. Anatomical ISC was calculated from the untransformed test responses and therefore depended only on the number of test repetitions.

