## Supplementary Materials for "Hyperalignment Reveals Shared Information in Idiosyncratic Fine-Grained Movie fMRI Patterns in Macaques"

### Supplementary Figures

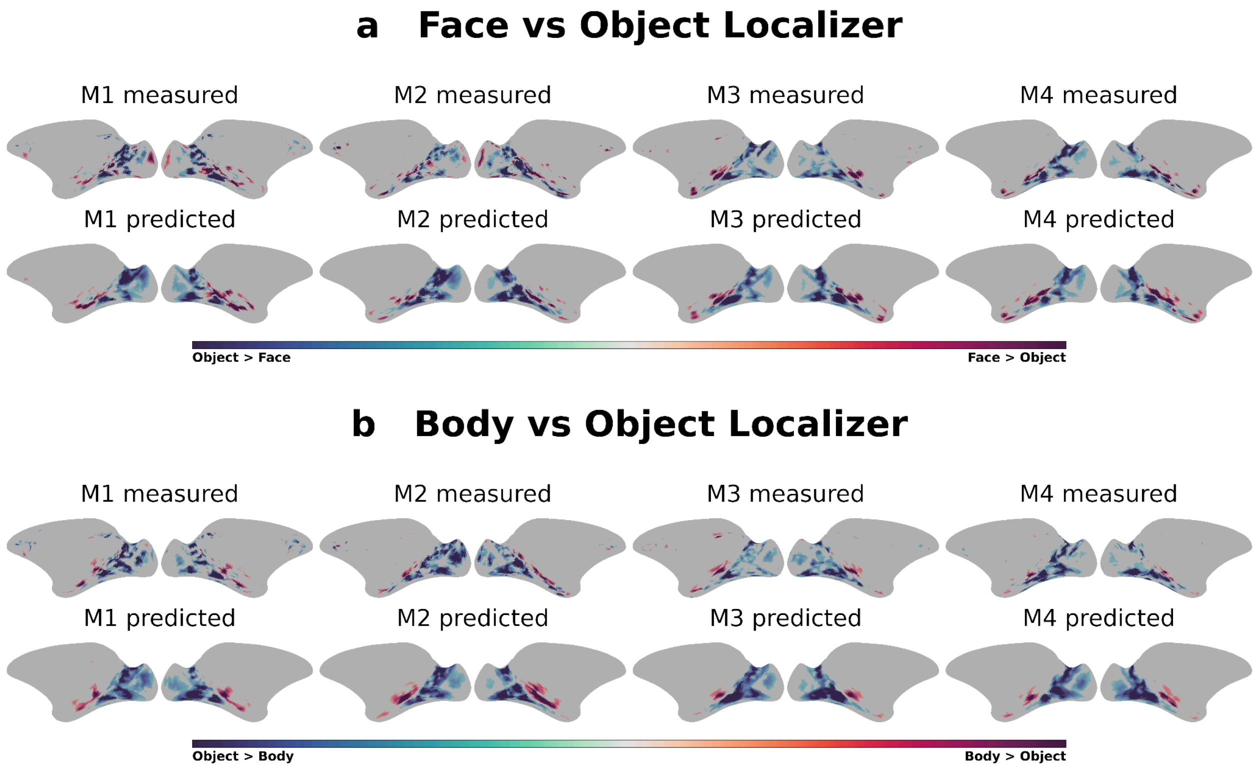


**Supplementary Figure S1: Hyperalignment-derived predictions recover individual face- and body-selective topographies in macaques.**
(a) Measured and predicted face-versus-object localizer maps for all four macaques (M1–M4).
(b) Measured and predicted body-versus-object localizer maps for all four macaques (M1–M4).
Color bars show t values.

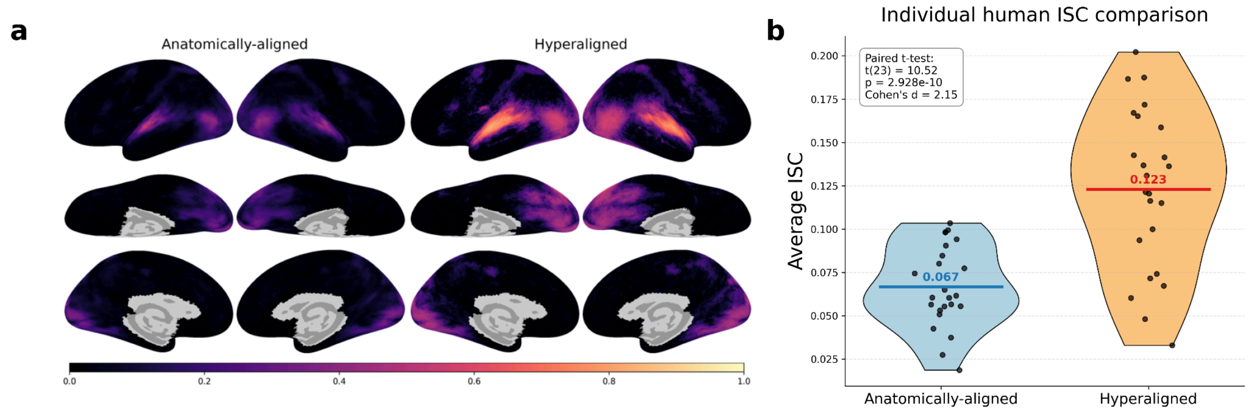


**Supplementary Figure S2: Hyperalignment increases inter-subject correlation of movie-evoked responses in humans.** (a) Group-average inter-subject correlation (ISC) maps for anatomically-aligned and hyperaligned human fMRI data during movie viewing. (b)

Distribution of average whole-brain ISC values across individual human subjects for anatomically-aligned and hyperaligned data. Each dot represents one subject, and horizontal lines indicate group means. Mean ISC increased from 0.067 under anatomical alignment to 0.123 after hyperalignment. A paired t-test showed a significant improvement after hyperalignment ($t(23) = 10.52$, $p = 2.928 \times 10^{-10}$, Cohen's $d = 2.15$).

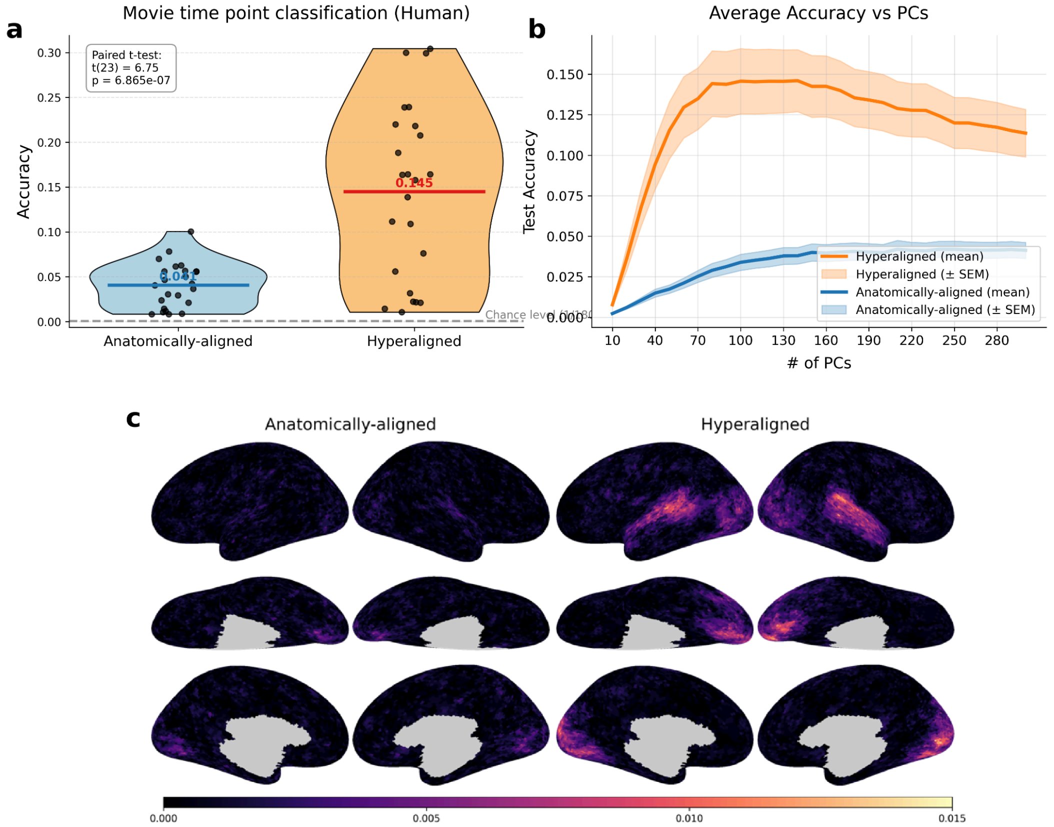


**Supplementary Figure S3: Hyperalignment improves between-subject movie time-point classification in humans.** (a) Whole-brain movie time-point classification accuracy for anatomically-aligned and hyperaligned human data. Each point denotes one subject, and horizontal lines mark group means. Hyperalignment increased mean accuracy from 0.041 to 0.145 (paired t-test: $t(23) = 6.75$, $p = 6.865 \times 10^{-7}$). The dashed line shows chance performance (1/1800). (b) Average classification accuracy across different numbers of retained principal components. Shaded regions denote ± SEM. (c) Searchlight-based classification accuracy maps for anatomically-aligned and hyperaligned data.

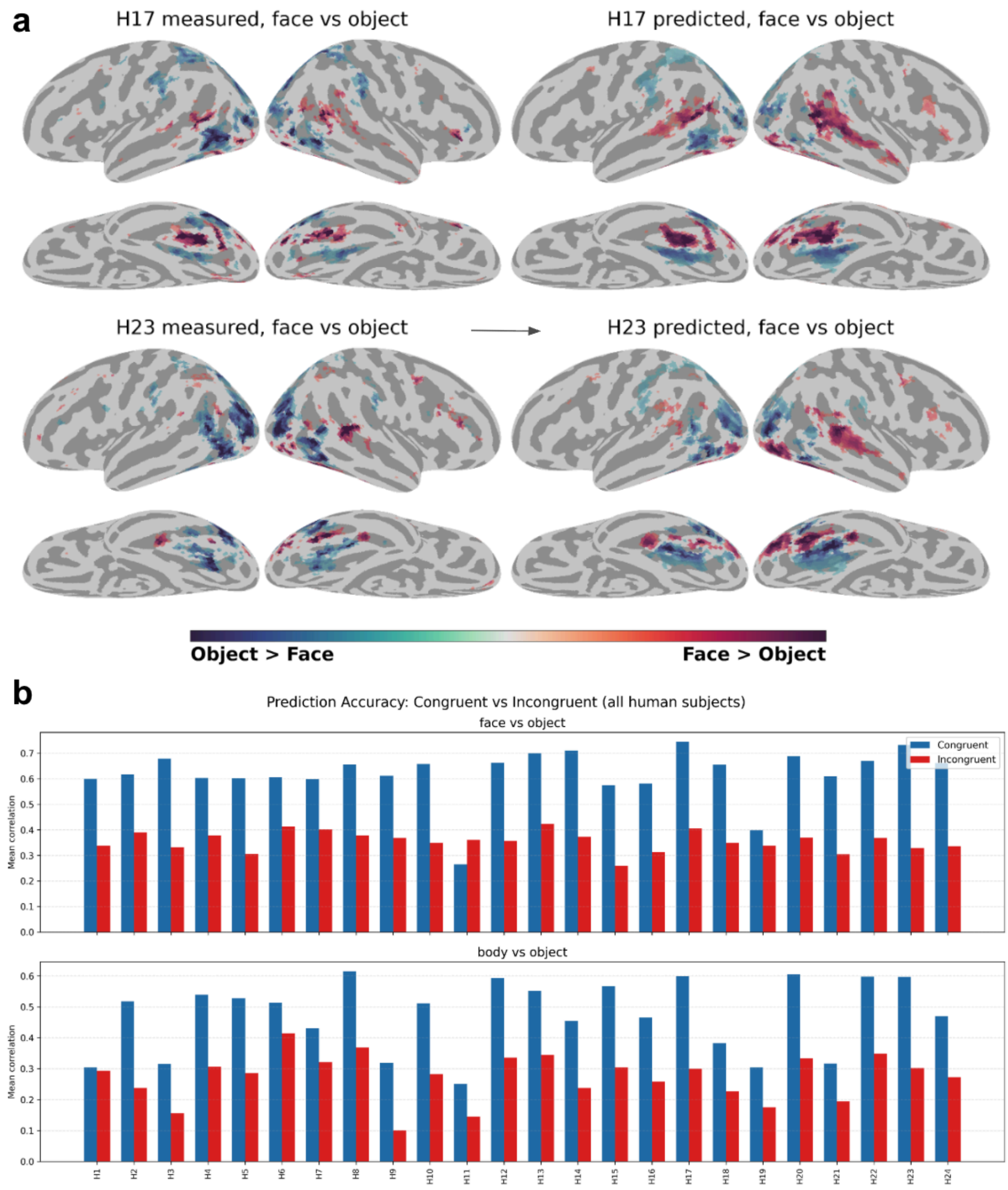


**Supplementary Figure S4: Hyperalignment enables prediction of individual functional topographies in humans.** (a) Measured and predicted face-versus-object localizer maps for two example human subjects (H17 and H23) on the cortical surface (full individual breakdown shown in Supp. Fig S5 and S6). Predicted maps accurately recovered subject-specific functional topographies. (b) Mean correlations between measured patterns and congruent (blue) or incongruent (red) predicted patterns for face-versus-object (top) and body-versus-object (bottom) localizers across 24 human subjects (H1–H24).

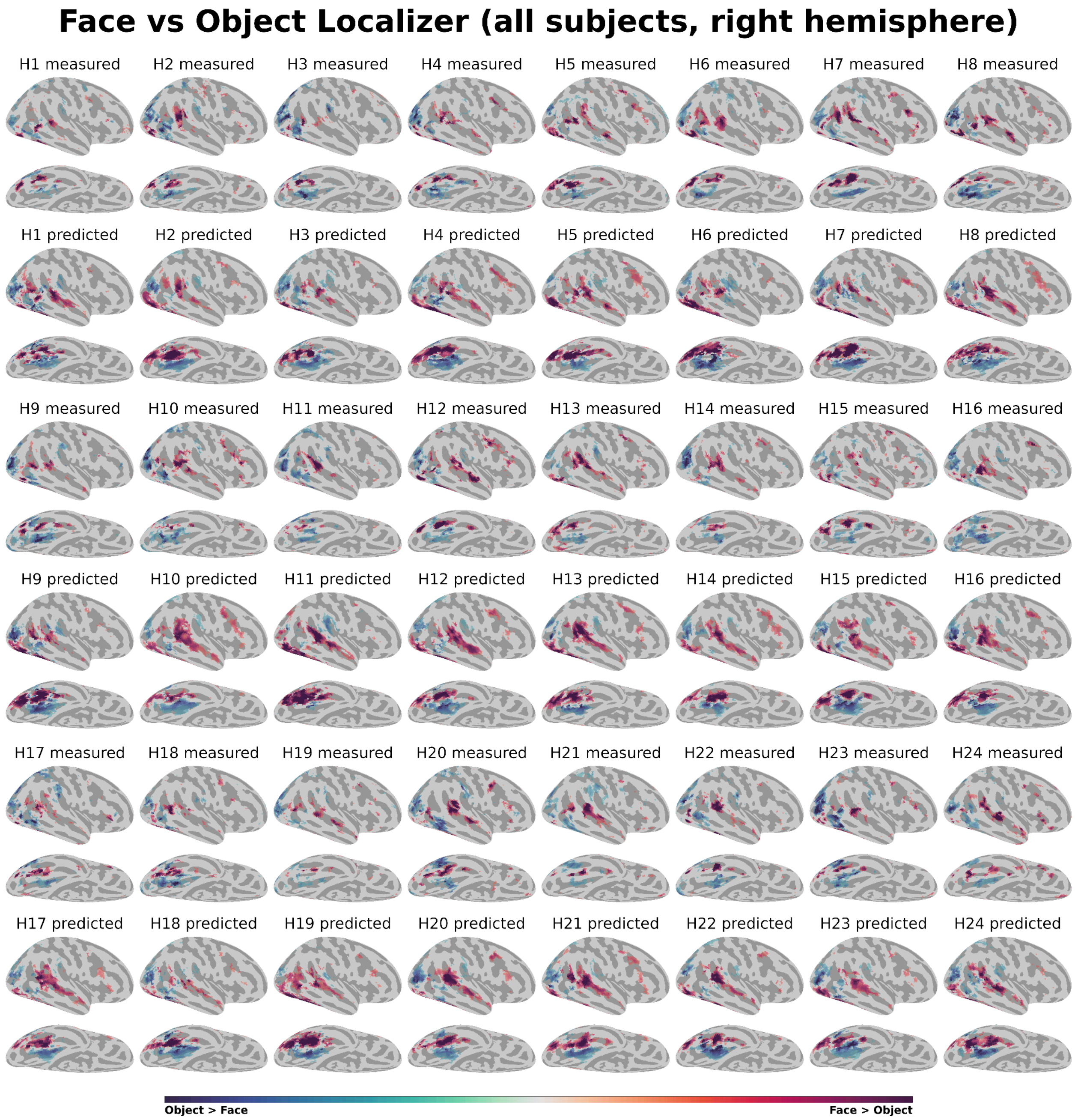


**Supplementary Figure S5: Measured and predicted face-versus-object localizer maps across all human subjects (right hemisphere only).** Individual measured and predicted face-versus-object localizer maps on the cortical surface for all 24 human subjects (H1–H24).

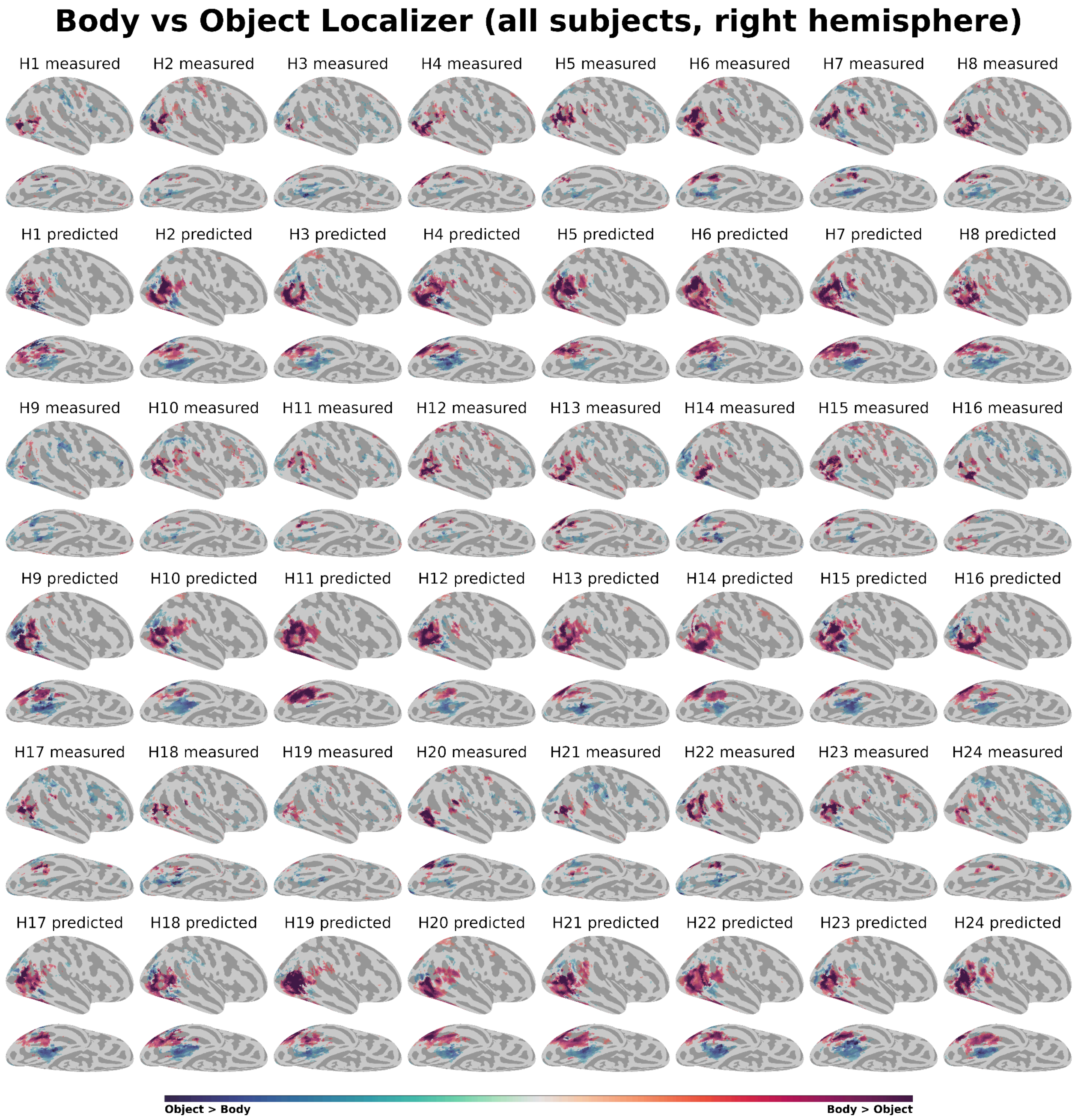


**Supplementary Figure S6: Measured and predicted body-versus-object localizer maps across all human subjects (right hemisphere only)**. Individual measured and predicted body-versus-object localizer maps on the cortical surface for all 24 human subjects (H1–H24).

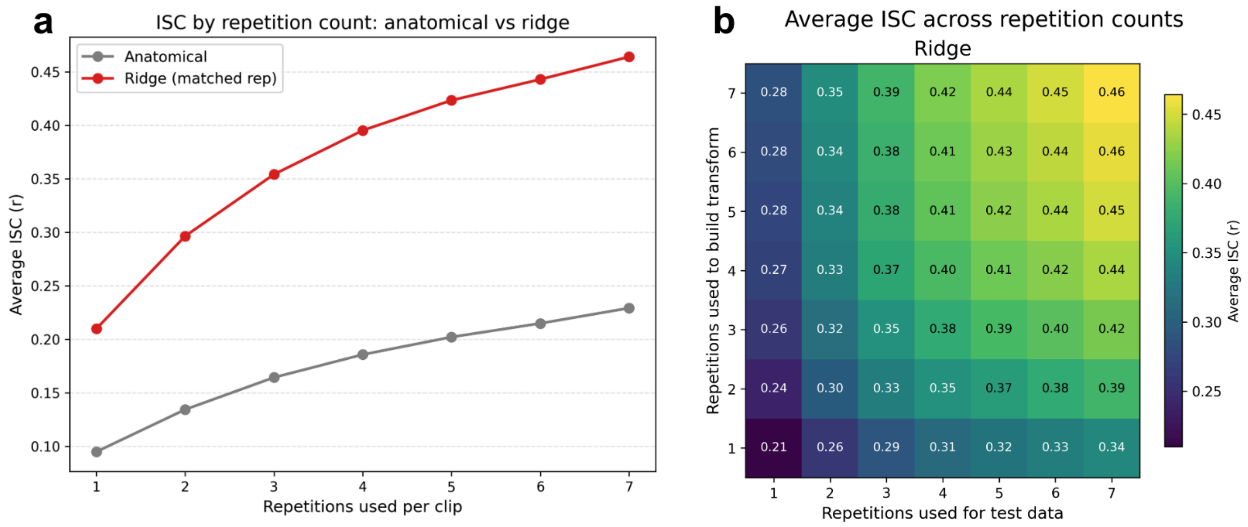


**Supplementary Figure S7: Effect of movie-repetition count on ISC after ridge hyperalignment.** Macaques viewed the same movie clips multiple times. (a) Mean whole-brain ISC for anatomical alignment and RHA when the same number of repetitions was used to estimate the ridge transformation and average the held-out test responses. Ridge hyperalignment using two repetitions exceeded anatomical alignment using seven repetitions. (b) Mean ISC for each combination of repetitions used to estimate the ridge transformation and repetitions used to average the held-out test responses. ISC improved as more repetition data were used, especially when more repetitions were averaged in the held-out test data.
